# A Comprehensive Phylogenetic and Bioinformatics Survey of Lectins in the Fungal kingdom

**DOI:** 10.1101/2021.04.01.438069

**Authors:** Annie Lebreton, François Bonnardel, Yu-Cheng Dai, Anne Imberty, Francis M. Martin, Frédérique Lisacek

## Abstract

Fungal lectins are a large family of glycan-binding proteins, with no enzymatic activity. They play fundamental biological roles in the interactions of fungi with their environment and are found in many different species throughout the fungal kingdom. In particular, their contribution to defence against feeders has been emphasized and extracellular lectins may be involved in the recognition of bacteria, fungal competitors and specific host plants. Their carbohydrate specificities and quaternary structures vary widely, but evidence for an evolutionary relationship within the different classes of lectins is provided by the high degree of amino acid sequence identity shared by the different fungal lectins. The UniLectin3D database contains 194 3D structures of fungal lectins, of which 129 are characterized with their carbohydrate ligand. UniLectin3D lectin classes from all origins were used to construct 107 lectin motifs in 26 folding configurations and to screen 1,223 species deposited in the genomic portal MycoCosm of the Joint Genome Institute. The resulting 33 485 protein sequences of putative lectins are organized in MycoLec, a publicly available and searchable database. The characterization of the lectin candidates in fungal genomes is based on systematic statistics regarding potential carbohydrate ligands, protein lengths, signal peptides, relative motif positions and amino acid compositions of fungal lectins. These results shed light on the evolution of the lectin gene families.

## Introduction

Fungi are unicellular or multicellular organisms found in terrestrial, marine and aquatic ecosystems. They adopt a wide range of ecological lifestyles (e.g., modes of nutrition) along the saprotrophism/mutualism/parasitism continuum (Naranjo-Ortiz and Gabaldón 2019; Richards et al. 2015; Stajich 2017). Saprotrophic, symbiotic and pathogenic fungal species compete or cooperate with multiple bacteria, fungal, plant or animal species. Specific signaling and developmental pathways are underlying the establishment and maintenance of the relationships with other organisms. Self/non-self discrimination is at the very core of the fungal hyphae development and involves surface-surface molecular interactions (Fischer and Glass 2019). Similarly, recognition of the animal or plant surfaces play a key role in the pathogenic or mutualistic fungal symbioses. These interactions rely on a wide range of secreted proteins, including lectins. Lectins are ubiquitous carbohydrate-binding proteins playing a crucial role in self/non self-recognition through specific interactions with complex glycans present on the surface of proteins, microorganisms or tissues (Lis and Sharon 1998). These small proteins are therefore involved in a wide range of biological processes such as reproduction and development, the establishment of biotrophic associations, but also in many pathogenesis-related mechanisms, such as host-pathogen interactions and tumor metastasis. A large number of lectins have been identified in filamentous fungi and yeasts (Goldstein and Winter 2007; Guillot and Konska 1997; Nordbring-Herz and Chet 1986; Pemberton 1994; Singh, Bhari, and Kaur 2011). They are generally considered as defence-related proteins. This role is well documented in mushroom-forming fungi where lectins are known to protect these reproductive structures from hyphal grazers, such as nematodes, slugs, snails and insects, through their nematotoxic and entomotoxic activities (Sabotič, Ohm, and Künzler 2016).

Except for their role in defence, the current body of information on the biological and ecological roles of lectins in saprotrophic, mutualistic and pathogenic fungi is scarce. Some yeasts produce flocculins that play a role in the formation of multicellular structures of interest for brewers and winemakers (Goossens and Willaert 2010; Reynolds 2018). Pathogenic yeasts, such as *Candida albicans*, rely on adhesins to initiate the colonization of epithelial hosts (Willaert 2018), while filamentous pathogenic fungi, such as *Aspergillus fumigatus*, produce soluble lectins involved in host recognition and able to trigger signaling pathways leading to lung inflammation (Houser et al. 2013; Richard et al. 2018). Less is known about the role of lectins in the establishment of fungal-plant or fungal-animal mutualistic symbioses. A role of fungal lectins in the establishment of the mutualistic ectomycorrhizal symbiosis between the basidiomycete *Lactarius deterrimus* and its host tree, *Picea abies*, has been suggested (Giollant et al. 1993) but not yet confirmed by genetic disruption of the corresponding genes. A galactose-specific galectin acts as a chemoattractant for the recruitment and adhesion of the cyanobacterial *Nostoc* symbiont to the hyphae of *Peltigera canina* leading to a lichen symbiosis (Díaz et al. 2011). The involvement of fungal lectin has since been established for a large number of lichen associations (Singh and Walia 2014).

The paucity of knowledge on the biochemistry and role of fungal lectins is reflected by the lowquality annotations of lectins in international protein and genome databases. Moreover, a limited number of 3D structures of fungal lectins is currently available. A recent review reported 26 different fungal lectins from ten different folds for 100 crystal structures (Varrot, Basheer, and Imberty 2013). Using a classification based on 3D X-ray structures and protein sequences, the curated UniLectin3D database (www.unilectin.eu/unilectin3D) (Bonnardel et al. 2019) can now be used to generate conserved sequence motifs based on known lectin classes. Those conserved motifs can further be used to characterize the lectin repertoire, so-called lectomes, of a given species. This lectin repertoire is indexed in LectomeXplore (Bonnardel et al. 2021). The objective of this study was to develop a novel approach for efficiently and accurately predicting and classifying lectin proteins in Fungi. We have used UniLectin3D-based motifs to predict lectins in published fungal genomes available at the Joint Genome Institute (JGI) MycoCosm genomics portal (http://jgi.doe.gov/fungi) (Grigoriev et al. 2014). This database aims to provide access of genomes representative of most fungal lifestyles and supports integration, analysis and dissemination of fungal genome sequences and other ‘omics’ data by providing interactive webbased tools (Grigoriev et al. 2014). At the time of this study in 2020 this database contained approx. 1500 genomes (from 1223 species). From these, the resulting 35 460 identified lectin domains on 33 485 sequences of putative lectins can be searched and analyzed on the MycoLec web site (https://www.unilectin.eu/mycolec/). In order to identify possible relationships between the lectin repertoire and the diverse ecological niches of fungal species, we focused our survey on the Agaricomycetes (Basidiomycota phylum). The rationale is the availability of a large set of genomes and transcriptomes for this fungal lineage that covers species with contrasted ecologies along the saprotrophism/symbiosis continuum. In this study we used this large-scale processing to comprehensively map the different groups of lectin genes and explore the origin and evolution of these gene families in Fungi. Then, we focused on Agaricomycetes to single a few lectin candidates out of the initial thousands to be further characterised with functional analyses.

## Materials and Methods

### Analysis of the fungal lectin classes

Classification of lectins has been described before (Bonnardel et al. 2021). Briefly lectins available in UniLectin3D (Bonnardel et al. 2019) were used to extract the lectin domain sequences and lectins fold sharing more than 20 percent of sequence similarities were grouped in the same class. This classification is applied to 3D-structures of fungal lectins.

### Fungal genomes

All proteomes available on the MycoCosm platform (mycocosm.jgi.doe.gov) (Grigoriev et al. 2014; Kohler et al. 2015) in 2020 were downloaded, resulting in the gathering of 1223 fungal proteomes. The taxonomy of each strain was retrieved using ncbitax2lin (github.com/zyxue/ncbitax2lin). When missing, the taxonomy was manually curated using JGI (taxonomy.jgi-psf.org) and UniProt taxonomy (www.uniprot.org/taxonomy). The phylogenomic tree of the fungal strains was obtained through the MycoCosm platform (Spatafora et al. 2017). Ecological guild were gathered using FUNGuild (Nguyen et al. 2016) and expert ecological annotation was performed for the 125 Agaricomycetes strains available at the time of this study.

### Identification and scoring of candidate fungal lectins

For each class, the lectin 3D domain sequences from which histidine tags were removed, were aligned with MUSCLE (Edgar 2004). The obtained alignments were used to generate a Hidden Markov Model (HMM) for each domain. HMMSEARCH (Potter et al. 2018) was then run to identify lectins in the proteomic dataset. Results were uploaded in the MycoLec database, which provides direct web access to the identified lectin candidates. Lectins were further filtered by a score, corresponding to the sequence similarity between the reference lectin domain consensus sequence and the predicted lectin domain sequence **(Bonnardel et al. 2021)**. This score was also used to organize the identified lectin candidates on the web database.

### Transcriptomics

Results of RNAseq experiments were compared to differentiate gene expression between the control condition and contact with a compatible plant in 14 mycorrhizal strains pertaining to the genomic dataset **(Kohler et al. 2015; Martino et al. 2018; Miyauchi et al. 2020; Murat et al. 2018; Peter et al. 2016)**. Among the strains, three are ericoid mycorrhizae (ERM), two are orchids mycorrhizae (ORM) and the other are ectomycorrhizae (ECM) (Table S1). Genes were considered as differentially expressed during mycorrhization compared to control when a log2 fold change above 2 or below -2 and an adjusted p-value below 0.05 were observed.

## Results

### Structural classification of fungal lectins in UniLectin3D

The 2021 version of Unilectin3D database contains 549 distinct lectin sequences and 2,278 lectin 3D structures, including 1,456 with their interacting ligands (Bonnardel et al. 2019). Among these lectins, 194 structures belong to fungal species. The 3D structures of fungal lectins are partitioned in 12 different folds (for a total of 35 in the database) and 20 classes (for a total of 109 in the database) (Figure 1 and Figure S1). Except for the ß-prism III fold, fungal lectin folds are shared with other organisms. The most abundant lectin 3D structures in the surveyed fungi are the ß-sandwich/Con-A-like fold (24%), the ß-propeller fold (18%) and the ß-trefoil fold (17%). When looking at classes, some are very specific of fungi, for example fungal fruit body lectins. The most represented classes include galectin-like (18%), PA14 yeast adhesin (15%), AAL-like (12%), fungal fruit body lectin (11%) and *Boletus* and *Laetiporus* ß-trefoil lectin (7%) (Figure 1). While lectins are often forming multivalent protein complexes through monomer oligomerisation, fungal lectins generate multivalent binding sites through internal repeats, as in the case of ß-trefoils and ß-propellers (Notova et al. 2020).

**Figure 1.**
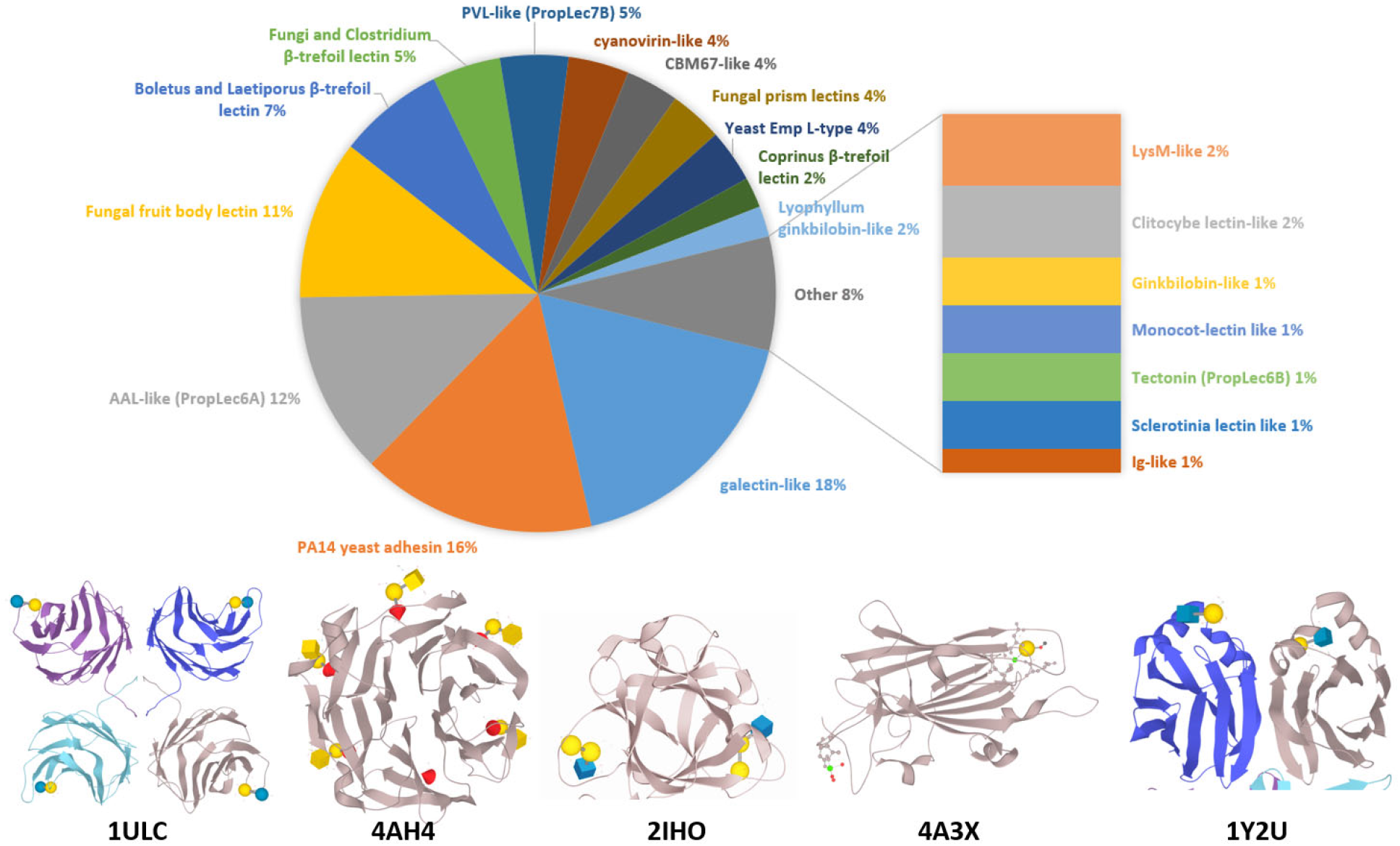
Distribution of fungal lectins 3D structures classes in UniLectin3D. Bottom: 3D view of most common folds for fungi lectins, using selected structures from the PDB (www.rcsb.org) (Burley et al. 2018): 1ULC, β-sandwich / ConA-like galectin-like from *Coprinopsis cinerea*; 4AH4, β-propeller AAL-like (PropLec6A) from *Aspergillus fumigatus*; 2IHO, β-trefoil / Fungi and Clostridium b-trefoil lectin from *Marasmius oreades*; 4A3X, β-sandwich / PA14 yeast adhesin from *Candida glabrata*; 1Y2U, β-sandwich / cytolysin-like Fungal fruit body lectin from *Agaricus bisporus*

### The MycoLec database of fungal lectomes

The MycoLec database (https://www.unilectin.eu/mycolec/) is built with the architecture of the LectomeXplore database. Its particularity is to be focused on Fungi. The prediction from the 1419 fungal genomes of the Mycocosm portal (February 2020 release) results in approximately 28 700 sequences of putative lectins for a score over 0.25 (Figure 2). These can be searched in multiple ways with queries based on the taxonomy, the lectin class or the protein name. Each candidate lectin is associated with sequence information including an alignment with the reference sequence of the predicted lectin class

**Figure 2:**
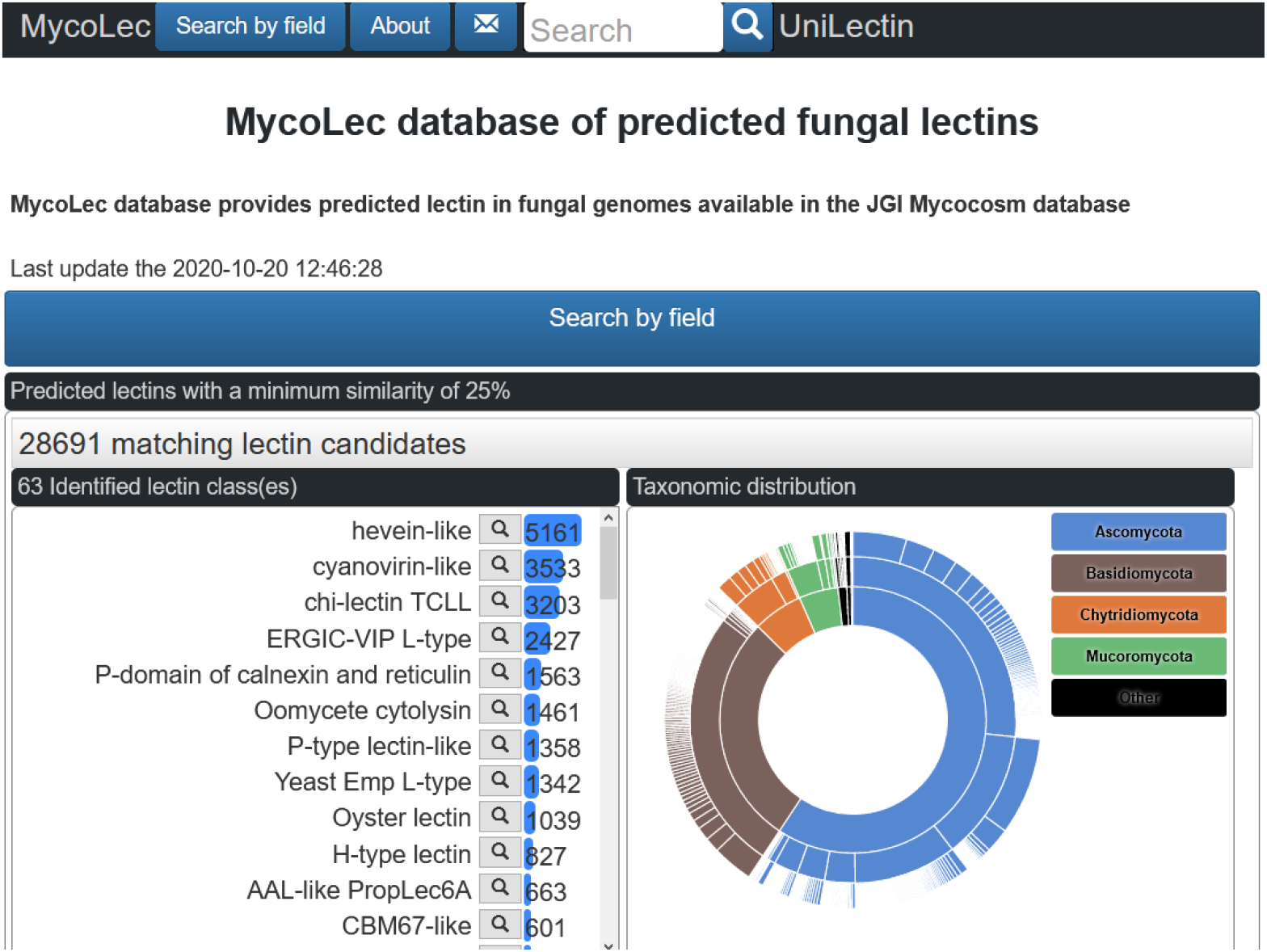
Access page of the MycoLec database with information about the distribution of fungal lectins as a function of classes and as a function of the fungal taxonomy.

### Identification of lectin domains in fungal proteomes

The MycoLec database contains 33,518 putative lectin sequences (28 691 with filter at 0.25 quality score) distributed across 27 folds and 63 lectin classes (Figure 3 and Figure S2). Notably, lectins have been found in all fungal genomes. 43% (12290) of the predicted lectins belong to a lectin class having a fungal origin in Unilectin3D (i.e. created using fungal proteins but not restricted to it). Lectins predicted using lectin motifs from fungal origins (Figure 3, purple) have, as expected, higher predicted scores, with the exception of the LysM-like lectins. Nevertheless, lectins are also predicted using lectin classes of non-fungal origins in Unilectin3D (or in the annotation). Surprisingly, Shiga-like toxin, Oomycete cytolysin and trefoil factors are predicted with globally high scores for lectins of other origins.

**Figure 3:**
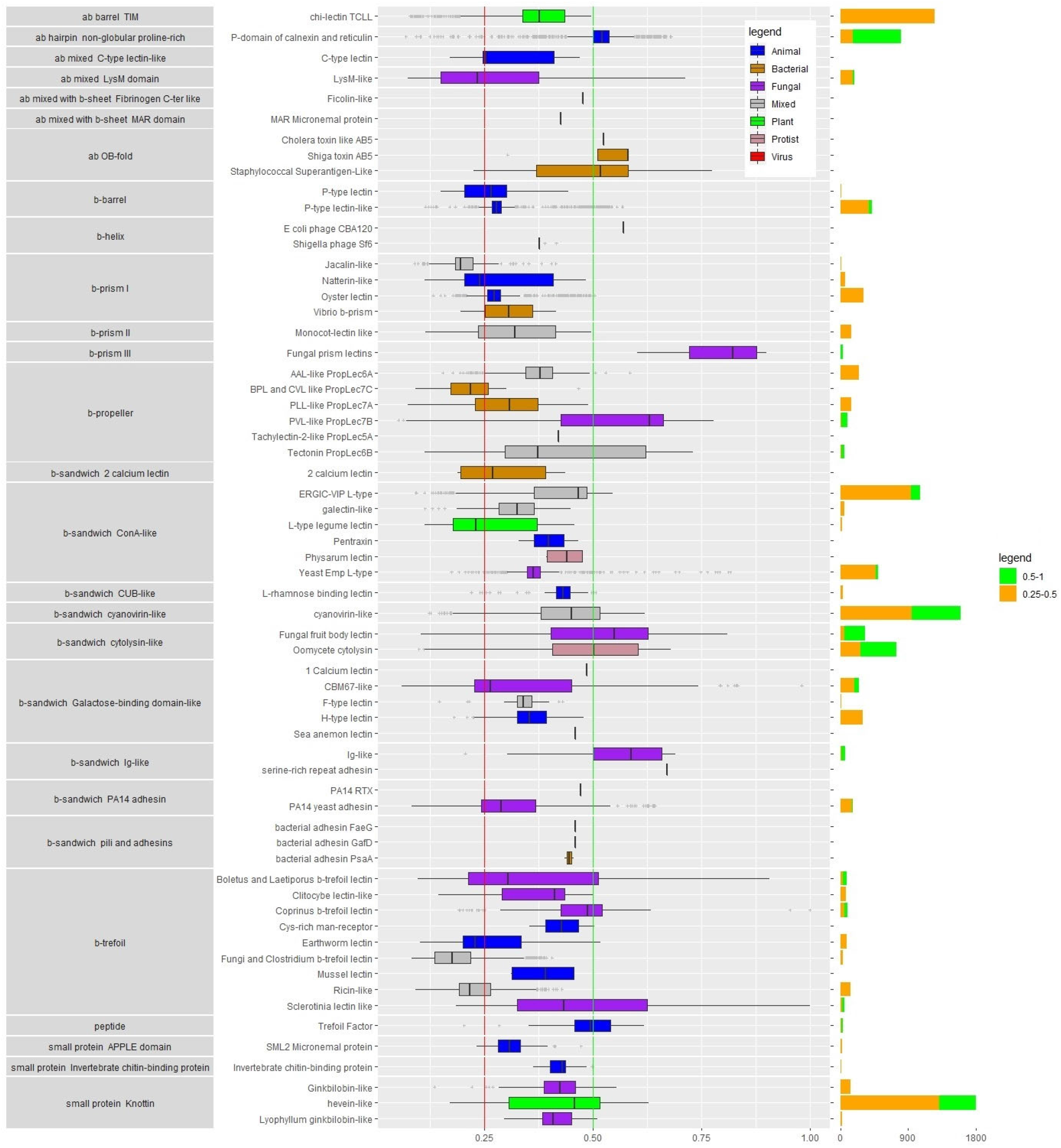
Distribution of the predicted fungal lectins sorted according to their fold and class. Boxplots are colored according to the structural origin of the lectin class defined by 3D structures. The right panel barchart represents the lectin distribution by class where green represents high confidence (score > 0.5) and orange, medium confidence predictions (score > 0.25).

### Phylogenetic distribution of fungal lectins

When looking at the lectome composition across the whole fungal kingdom, strong variations of lectin catalogs are observed depending on the fungal lineage (Figure 4 and Figure S3), with Agaricomycetes displaying the largest variety. The ubiquitous lectins are in general housekeeping proteins, involved in quality control of glycoproteins. This is the case of calnexin and calreticulin, acting as chaperones by selective binding of a glucose residue on misfolded glycoproteins and recruitment of folding factors (Oliver et al. 1997). The P-type like lectin is involved in misfolded glycoprotein ER-associated degradation (ERAD) in the cytosol. The carbohydrate recognition domain of p58/ERGIC-53 participates in glycoprotein export from the endoplasmic reticulum (Velloso et al. 2002). Other lectins are involved in non-self recognition, such as TCLL which is a degenerated chitinase (chitinase has an essential role in fungi), involved in antiviral activity. The oyster lectin also has an N-glycan (HMTGs) binding activity with microbicide activity.

**Figure 4.**
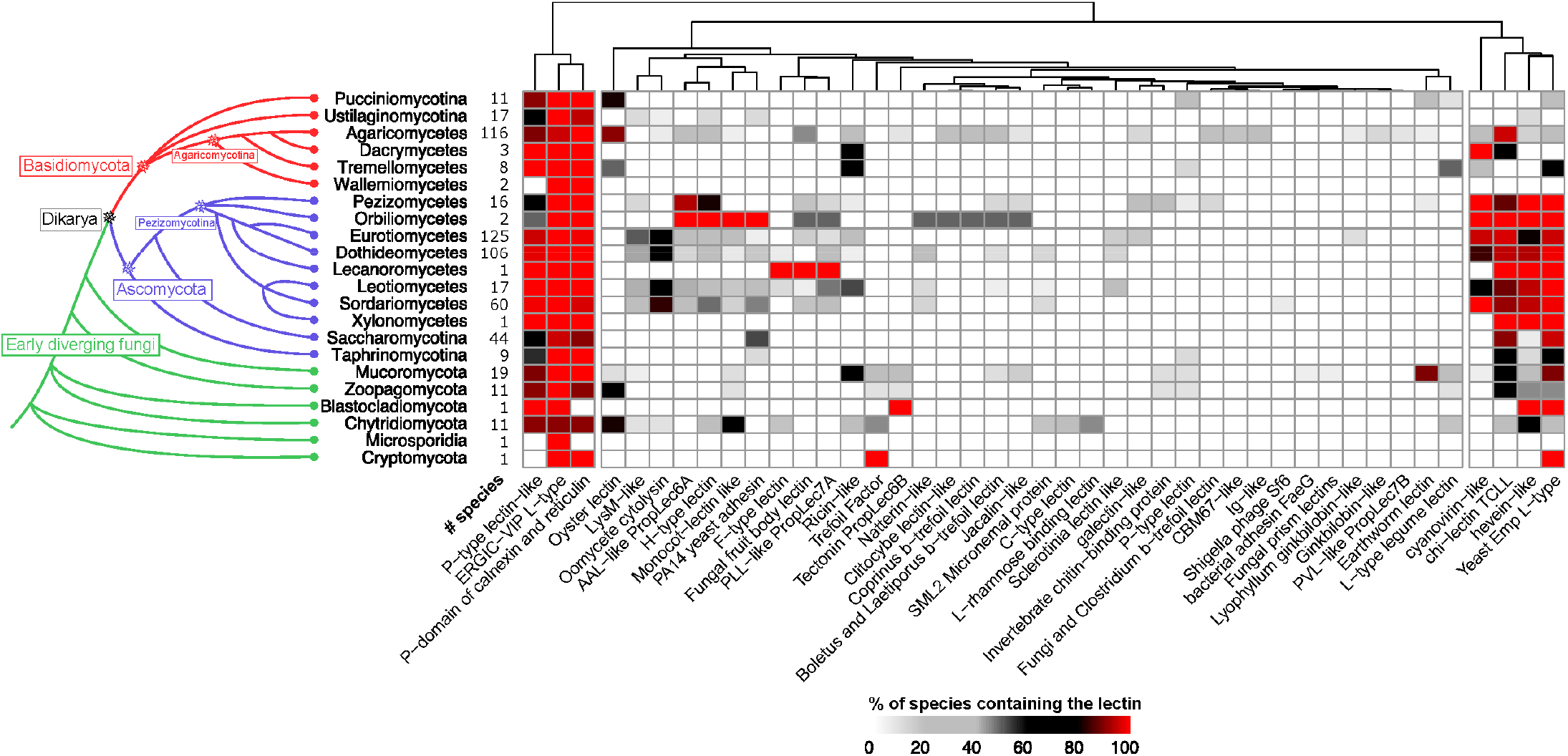
Distribution of predicted lectins by phyla and classes in MycoCosm genomes. Left, a phylogenetic tree showing the fungal divisions and classes. Right, Clustering of lectin classes found in 582 species. Lectins with a similarity score > 25% were used to compute the presence and abundance of the different lectin classes found in the MycoCosm genomes. The heatmap depicts lectin counts in each of the lectin classes, according to the colour scale displayed at the bottom of the figure.

Some lectin classes such as cyanovirin-like and fungal fruit body lectins were frequently identified within Agaricomycetes species. These families also show an expansion in specific species and genus suggesting that the corresponding lectins undergo rapid evolution.

The “house-keeping” lectins involved in quality control in the biosynthesis of glycoproteins are identified in all fungal classes, except Cryptomycota and Microsporidia. Cyanovirin-like, chi-lectin TCLL and Hevein and Yeast Emp L-type lectins are enriched in Ascomycetes and Blastocladiomycetes, but less present in comparison in Basidiomycetes. H-type lectin and monocot lectin-like are more present in Orbiliomycetes (containing nematode-trapping fungi), while the Lecanoromycetes (lichen fungi) are enriched in F-type and Fruiting body lectins. Chytridiomycota genomes stand out by a striking expansion of hevein lectins

### Distribution of lectin classes according to the nutrition modes

As observed above, Agaricomycetes (Basidiomycota) display the most diverse lectin composition. Since these fungi adopt a large variety of lifestyles, their lectomes were analyzed as a function of ecology. After filtering out the ubiquitous lectins, and those that are not present in Agaricomycetes (or as a single copy), the distribution of the 27 classes of interest was analyzed (Figure 5 and Table S2). Half of the species displays three of less classes of lectins (48/107) but five of them have more than 10 classes. The larger variety of lectin classes is observed in litter decayers with more than 10 different classes for some species, spanning more than 1000 genes coding for putative lectins. The wood decayers also display a larger variety of lectins than the pathogenic or mycorrhizal fungi. Endophyte shows the smallest number of lectins, although the sample is very limited in this study.

**Figure 5:**
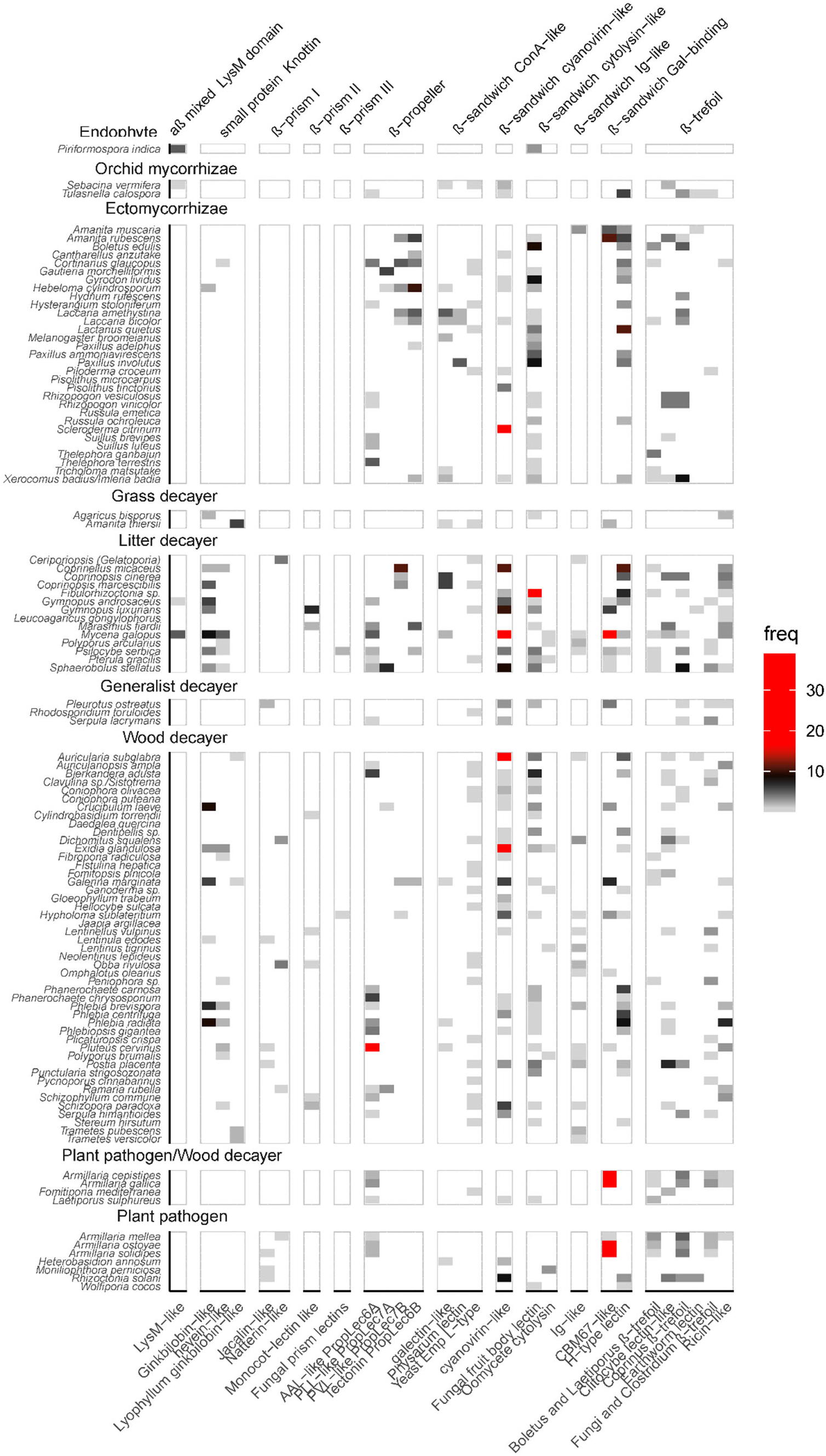
Lectomes, identified set of lectins in Agaricomycete species for a score higher than 25% of similarity to the reference. Species were grouped by ecology, and colors correspond to the number of lectins identified. Ubiquitous lectins and those present in only one species were filtered out.

The lectomes present large variation among Agaricomycetes species with no apparent correlation with the nutrition modes. The fungal fruit body lectins are prevalent, occurring in 50% of the investigated genomes. Cyanovirin-like proteins, 6-blades β-propellers and several β-trefoil lectin classes such as ricin-like and Coprinus-type and also present in many species. Cyanovirin and β-trefoil are widely distributed in eukaryotes and prokaryotes, while fungal fruit body lectins, also referred as actinoporin-type lectins (Sabotič, Ohm, and Künzler 2016) are highly specific to fungi, in both Ascomycetes and Basidiomycetes, with the only other occurrence in primitive plants such as Bryophyta and Hepatophyta.

Our study results in the identification of a new class, not yet characterised experimentally in fungi: the GalNAc-specific H-type lectin. This class is involved in self/non-self recognition in invertebrates, and has been first structurally characterized in snail eggs (Sanchez et al. 2006), but has never been isolated in fungi. A recent review on H-type lectins identified the corresponding sequence in the genome of several Agaricomycetes, including mycorrhizae *Tulaneslla calosporra* and wood decayer *Exidia glandulosa* (Pietrzyk-Brzezinska and Bujacz 2020). Our study demonstrates that a 30% occurrence in the investigated genomes. Similarly, a recent study reported the presence of the jacalin-like lectin in *Grifola frondosa* (syn. *Polyporus frondosus*) (Nagata et al. 2005), and the present work confirms the occurrence of this lectin in several other Agaricomycetes.

### Prediction of the lectome of Laccaria bicolor

*Laccaria* sp. are fungi pertaining to the Basidiomycota phylum and the Agaricomycetes Class. It forms ECM with a wide range of trees. *L. bicolor* is used as a model for ECM studies. Only one strain for *L. bicolor* is publicly accessible at the time of this study and so integrated in our dataset. A search with a 0.25 cutoff identifies 13 different classes of lectins in the genome (Table S3).

Filtering out house-keeping lectins and poorly scored predictions leaves eight lectins that are displayed in Figure 6. Two of them are proteins found in many organisms: Cyanovirin and Oyster lectin. Cyanovyrin, originally from cyanobacteria (Bewley 2001) has also been identified in plants and fungi, but its function is not elucidated. Oyster lectin has been recently found in bivalves (Unno et al. 2016) and is predicted to occur in many organisms, where its mannose-binding function is not yet clarified. The six other lectins are well characterized in different fungi where they play a role in defence against pathogens. Tectonin, structurally characterized in *L. bicolor* (Sommer et al. 2018), binds to methylated sugars that are present in nematodes (Wohlschlager et al. 2014), while the galectin-like lectin CGL2 from *Coprinopsis cinerea* inhibits the development of nematodes (Butschi et al. 2010). Interestingly, the *L. bicolor* lectome contains a full panel of fungi-related lectins. *L. bicolor* is a model for studying the establishment of ECM with the *Populus tremula x alba* tree (Martin and Selosse 2008). It is therefore worth trying to find out whether its many lectins could be involved in this process. Transcriptional regulation of lectin genes has been investigated in 14 mycorrhizal associations and RNAseq experiments were undertaken to differentiate gene expression with and without contact with a compatible plant **(Kohler et al. 2015; Martino et al. 2018; Miyauchi et al. 2020; Murat et al. 2018; Peter et al. 2016)** (Table S1). For most lectins, no variation in expression level was observed upon contact with the host plant, suggesting that only a restricted set of lectins might be involved in mycorrhization (Figure S4). Nevertheless, several *L. bicolor* lectins, i.e. tectonin, β-trefoil lectins, cyanovirin and fungal fruiting body lectins, are upregulated during the mycorrhization (Figure S5). However, this observation is not extended to the other fungi/tree association, and in some cases, lectin transcription is down-regulated during the mycorrhization. Nonetheless, the differential expression of lectins in fungi interacting with trees compared to the isolated condition may show the importance of specific lectins for some symbiotic-host interactions. Further studies will be necessary to analyze the role of lectins in the establishment of symbiosis.

**Figure 6:**
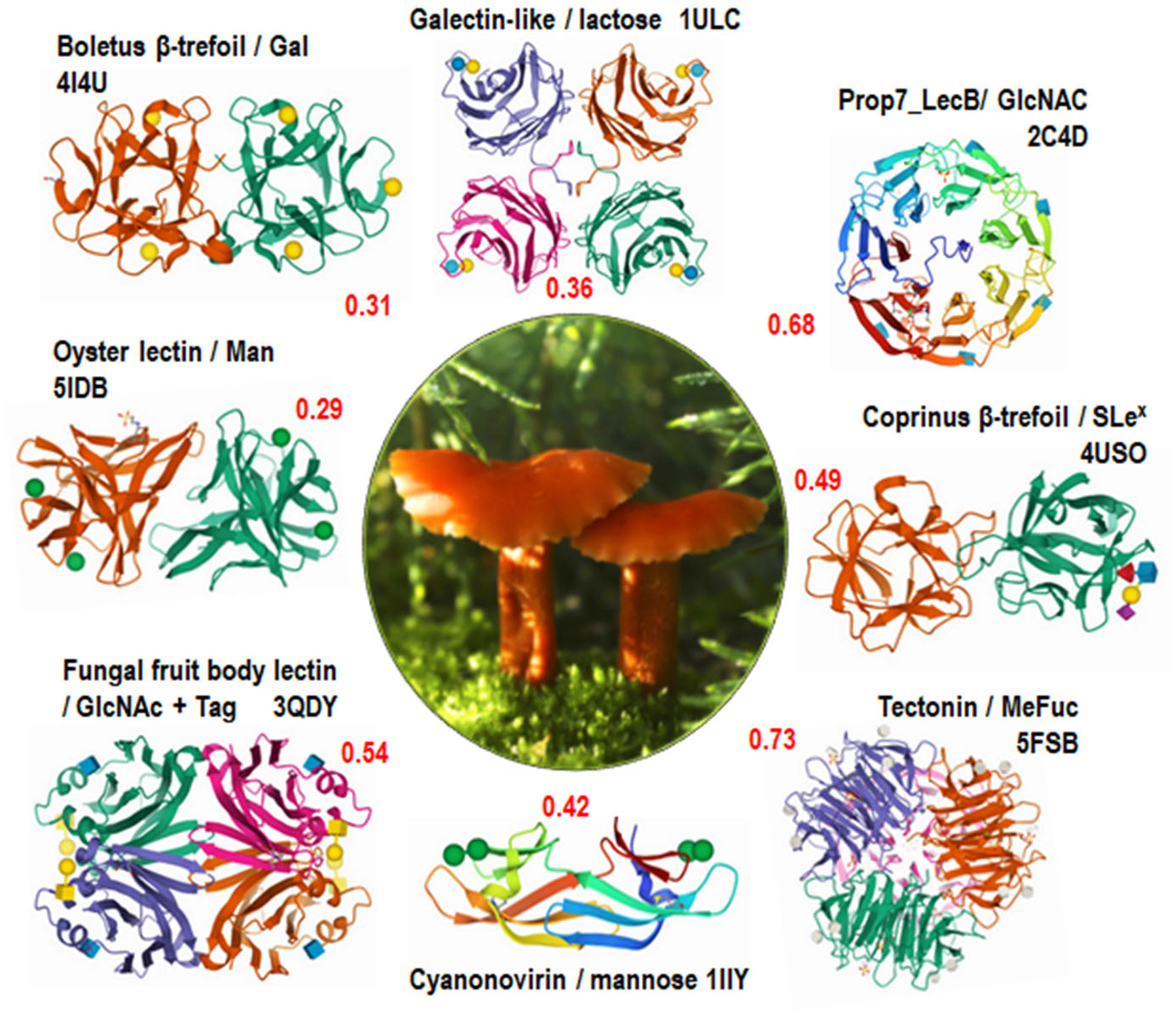
Part of the predicted lectome of *Laccaria bicolor* with scores indicated in red. Only lectin classes with strong score prediction and conserved binding sites have been selected and represented using Pymol (https://pymol.org/). Representative images for each class are from the PDB (www.rcsb.org) (Burley et al. 2018) and are generally from other species, except the tectonin which is from *L. bicolor*.

## Discussion and Conclusions

The aim of this project was to explore lectin composition and abundance across and in the context of the fungal kingdom. A large panel of 33,518 fungal lectins belonging to 63 distinct lectin classes have been identified using 1223 fungal strains from the MycoCosm database and can be accessed in an online interactive database available at unilectin.eu/mycolec. The MycoLec database can be searched and browsed to explore prediction results ranked by similarity score. Each predicted lectin is available together with the identified lectin domain in which the binding site can be compared to the reference motif. Significant differences have been shown to distinguish the fungal taxonomic classes. Ecological information was obtained for a part of those fungi allowing us analyze the putative relation of lectins with specific ecology.

Based on the Agaricomycete fungal class, we showed that lectomes vary with lifestyle and that saprophytic fungi, living on decaying wood and litters, have a larger variety of lectins than symbionts and pathogens. However, a statistical analysis is necessary to demonstrate the association of lectin with specific ecological traits. Previous analysis of glycoside hydrolases in the Agaricales order (Ruiz-Dueñas et al. 2020), led to reconstruct the history of this gene family and investigate the expansion and contraction event. Such a study seems to be challenging concerning the evolution of lectins because of the observed dispersion of genes within the dataset.

*Laccaria bicolor*, a model fungus for studying ECM, includes interesting predicted lectins with distinct folds and these proteins are up -regulated in the presence of the associated host plant and provide a lead for further analysis.

The present study brings a new perspective to the broad diversity of fungal lectomes. Even if a large number of fungal lectins were already identified, we provide the tools to better appreciate the extent of this repertoire. We could show that some lectin classes previously identified in invertebrates (H-lectins, oyster lectins) or in plants (jacalin, Ginkbilobin-like) turn out to be widely distributed in Agaricomycetes. Some of these new lectins could have useful application in biotechnology or as anti-viral compounds (El-Maradny et al. 2021) and the MycoLec database is readily available to be mined for novel lectins.

## Supplementary Materials

Figure S1: Distribution of lectin folds and classes of fungal lectin with 3D structures in Unilectin3D database, Figure S2: Distribution of folds (left) and classes (right) of predicted lectin sequences in MycoLect, Figure S3: Distribution of predicted lectins by species in MycoLect, Figure S4: Impact of mycorrhization of 14 fungal strains with their corresponding host plant on lectin expression, Figure S5: Differential expression of lectins within mycorrhizal fungi upon plants interaction, Table S1: Fungal species investigated in the exploration of the lectins induced and repressed during their mycorrhization with a compatible plant host, Table S2: Lectin content in the predicted proteomes of the Agaricomycetes fungal class sorted by ecological niche, Table S3: Details of lectins identified in the genome of *Laccaria bicolor*.

## Author Contributions

Conceptualization, A.I., F.L. and F.M.; methodology A.L. and F.B.; formal analysis A.L, F.B. and Y-C.D.; data curation A.I., A.L, F.L. and F.M.; writing— original draft preparation, A.L and F.B; writing—review and editing, A.I., F.L. and F.M.; visualization, A.L, F.B. and Y-C.D.; supervision, A.I., F.L. and F.M.; funding acquisition, A.I. and F.L. All authors have read and agreed to the published version of the manuscript.

## Funding

F.B. is funded by the ANR PIA Glyco@Alps (ANR-15-IDEX-02), A.L. is funded by a postdoctoral scholarship from the Beijing Advanced Innovation Center for Tree Breeding by Molecular Design, Beijing Forestry University, Beijing, China

## Acknowledgments

The authors acknowledge support by the ANR PIA Glyco@Alps (ANR-15-IDEX-02), the Alliance Campus Rhodanien Co-funds (http://campusrhodanien.unige-co-funds.ch), the Laboratory of Excellence ARBRE (ANR-11-LABX-0002-01), the Region Lorraine and the European Regional Development Fund.

## Conflicts of Interest

The authors declare no conflict of interest. The funders had no role in the design of the study; in the collection, analyses, or interpretation of data; in the writing of the manuscript, or in the decision to publish the results.

## Notes

https://www.unilectin.eu/mycolec/

### Competing Interest Statement

The authors have declared no competing interest.

