## Supplemental information for "A Comprehensive Phylogenetic and Bioinformatics Survey of Lectins in the Fungal kingdom"

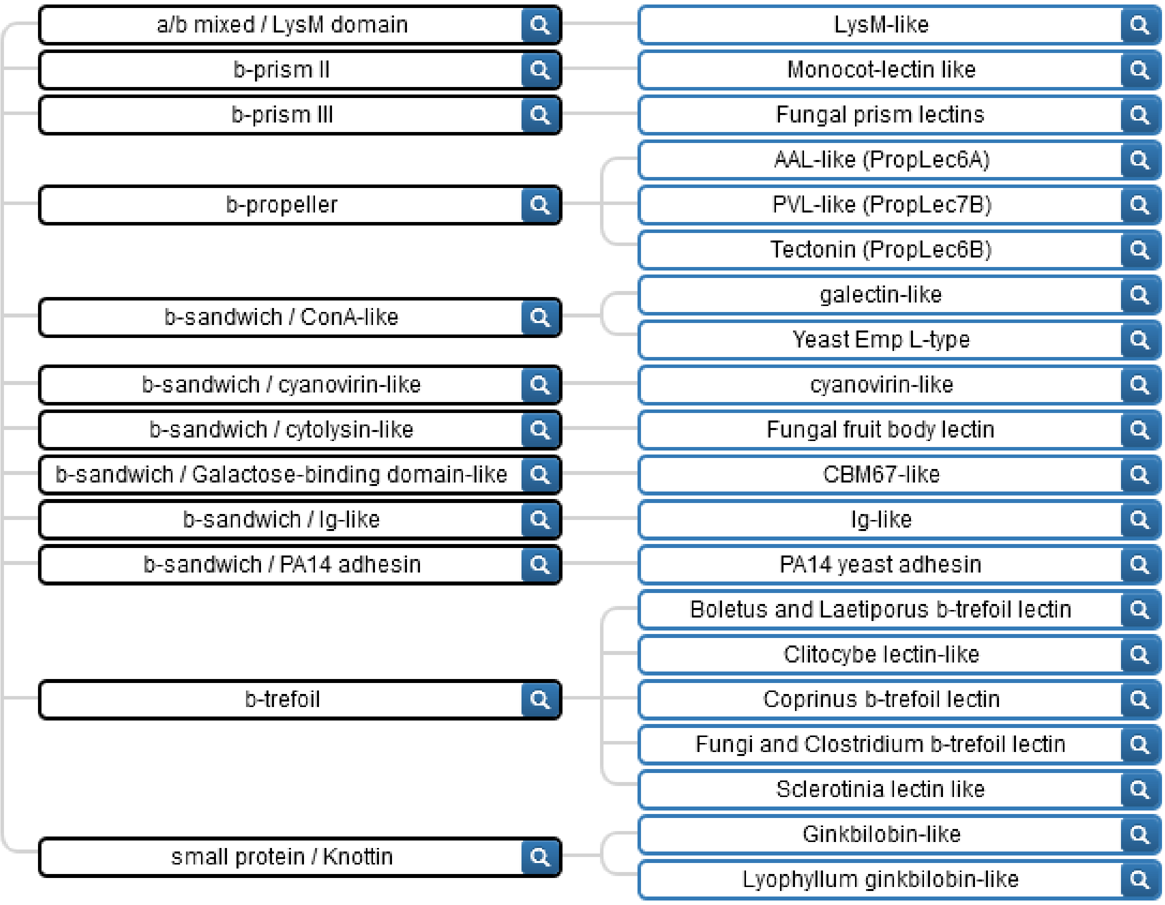


**Figure S1**: Distribution of lectin folds and classes of fungal lectin with 3D structures in Unilectin3D database.

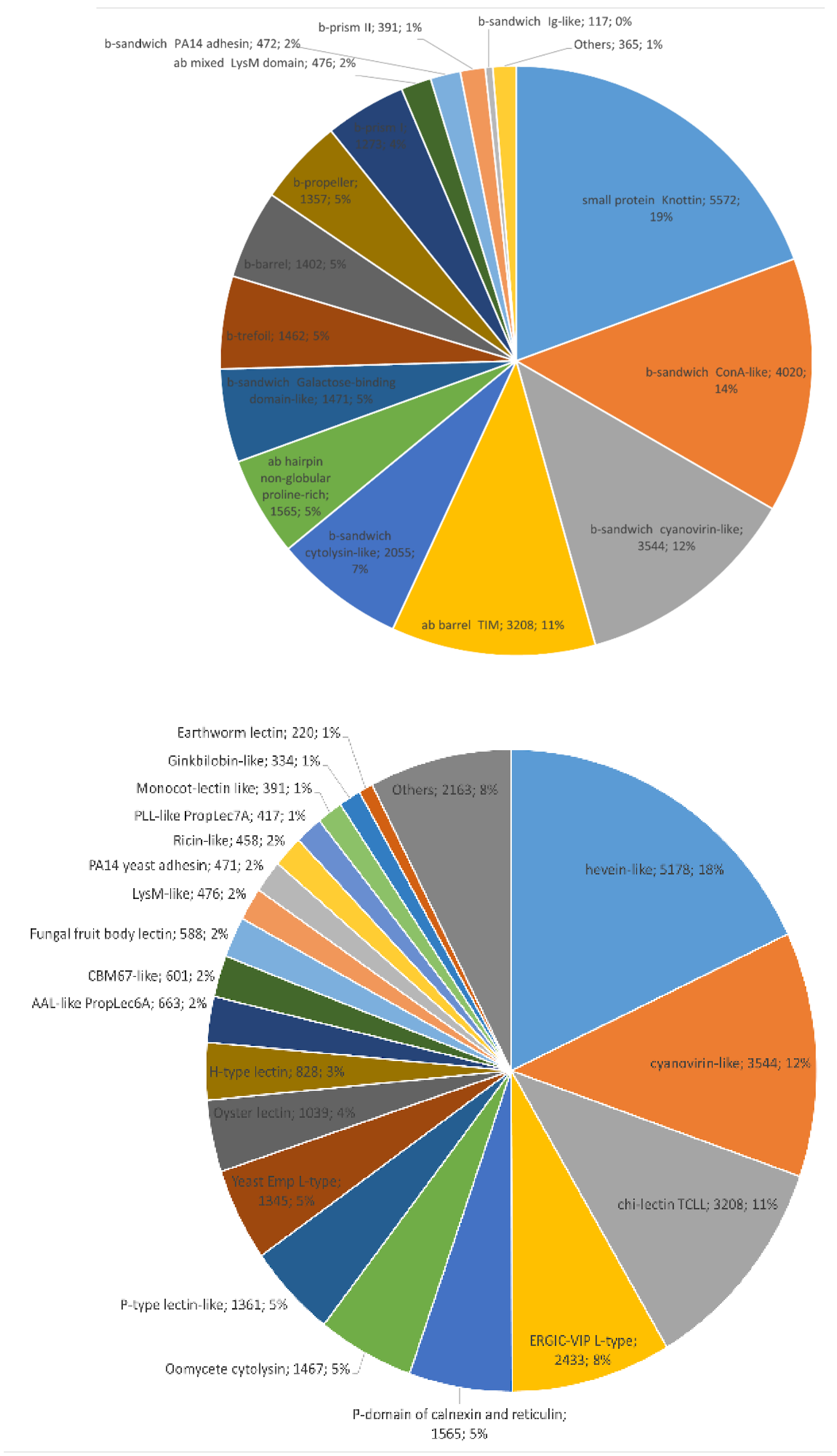


**Figure S2**: Distribution of folds (left) and classes (right) of predicted lectin sequences in MycoLec. Only lectin sequences with a similarity score > 25% were used.

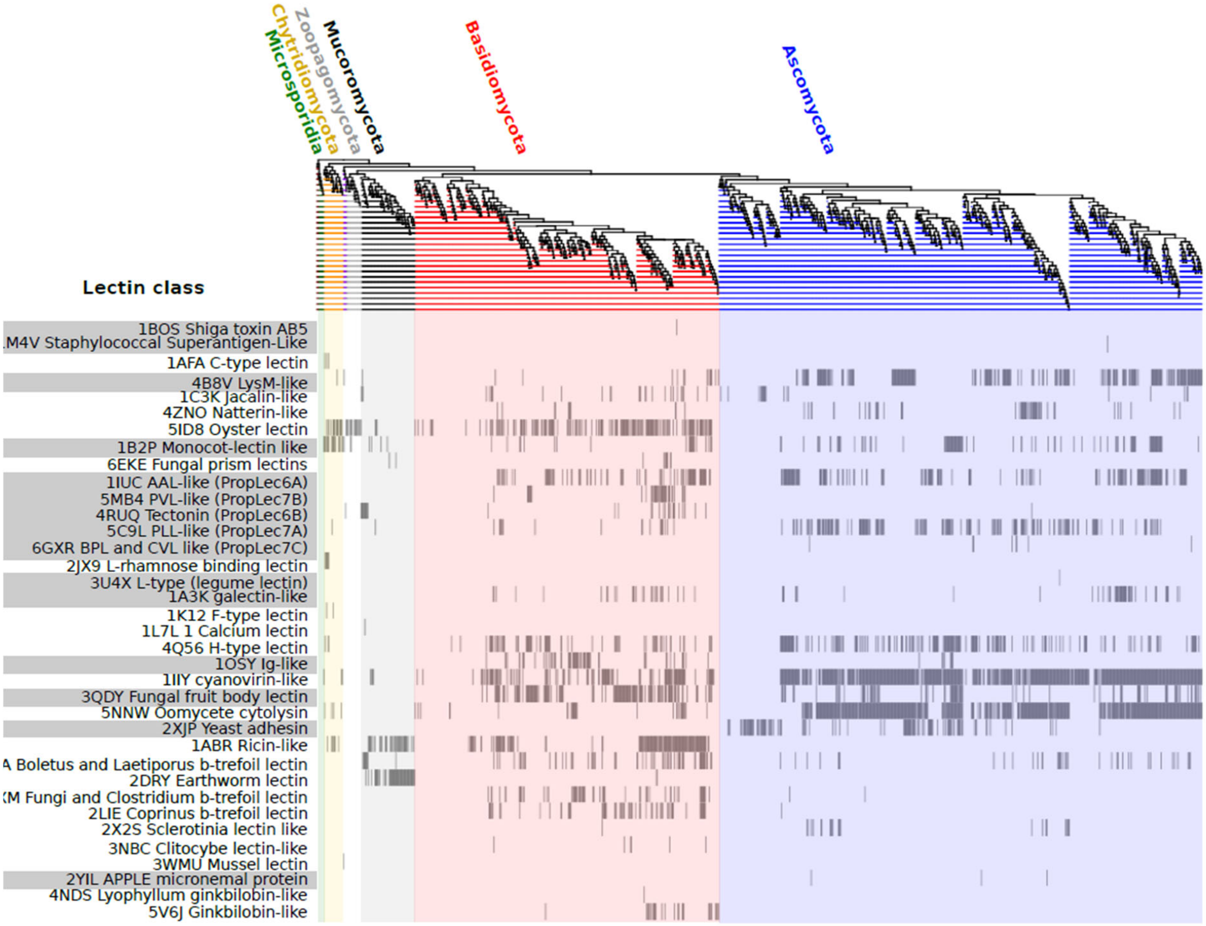


**Figure S3: Distribution of predicted lectins by species in MycoLec.** Each vertical line corresponds to a fungal strain organized according to their phylogenetic relationship as displayed by the tree. Left, Clustering of lectin classes. Lectins with a similarity score > 25% were used to detect the presence of the different lectin classes found in the MycoCosm genomes.

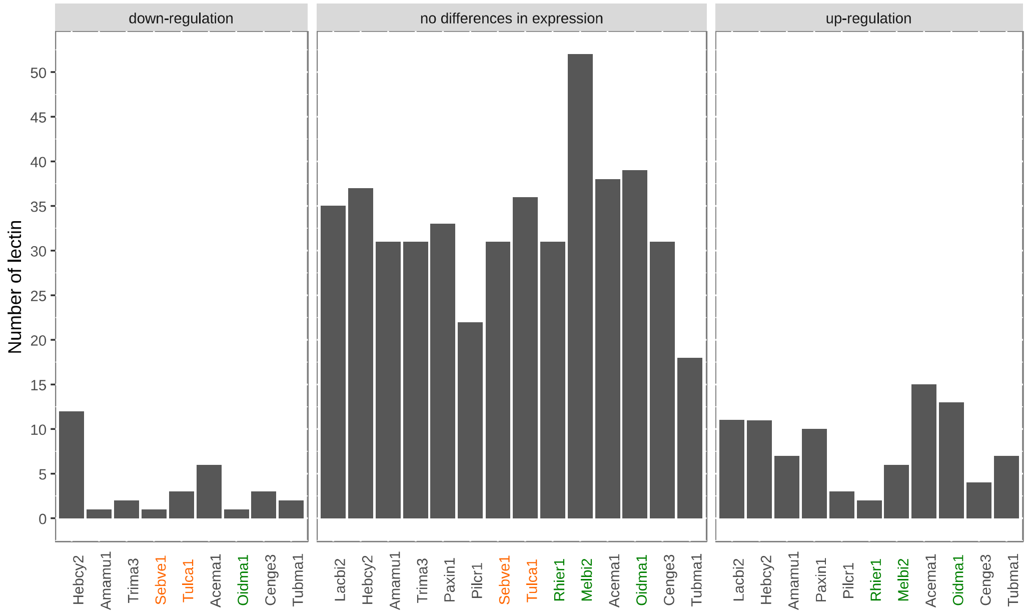


**Figure S4**: **Impact of mycorrhization of 14 fungal strains with their corresponding host plant on lectin expression.** Each bar corresponds to a strain annotated by a tag used in the MycoCosm database to refer to the specific strain and genomic assembly. Tags are colored according to the mycorrhizae type: grey ectomycorrhizae, green ericoid mycorrhizae, orange orchid mycorrhizae.

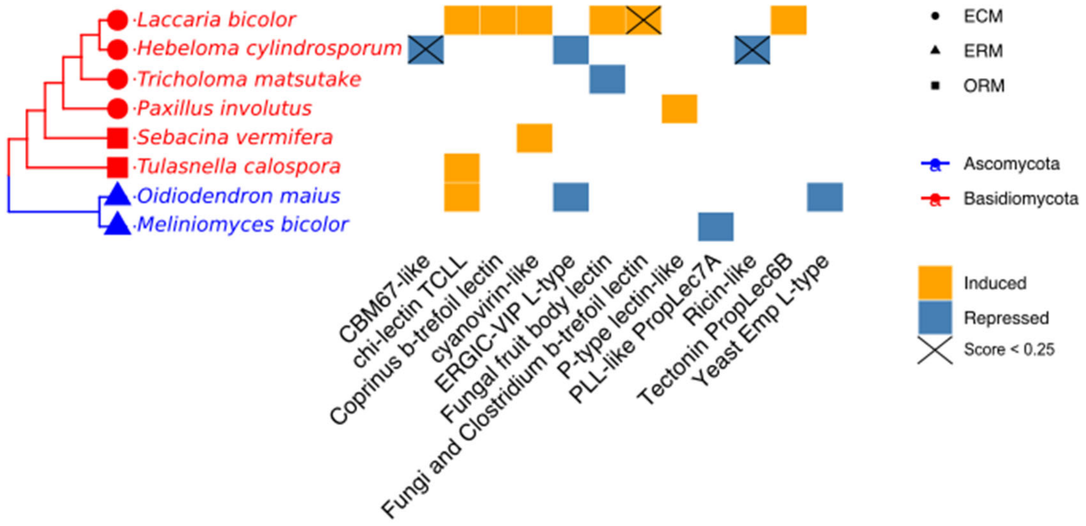


**Figure S5: Differential expression of lectins within mycorrhizal fungi upon plants interaction.** Lectins with invariable expression are not represented. Three species are ericoid mycorrhizae (ERM), two are orchids mycorrhizae (ORM) and the other are ectomycorrhizae (ECM).

**Table S1: Fungal species investigated in the exploration of the lectins induced and repressed during their mycorrhization with a compatible plant host.** ECM: ectomycorrhizae, ORM: orchids mycorrhizae, ERM: ericoid mycorrhizae

| JGI ID | Fungal species | Plant host | Analysis Method | Mycorrhizae type | Ref |
|---|---|---|---|---|---|
| Amamu1 | *Amanita muscaria* | *Populus tremula tremoloides* | CLC | ECM | 1 |
| Hebcy2 | *Hebeloma cylindrosporum* | *Pinus pinaster* | CLC | ECM | 1 |
| Paxin1 | *Paxillus involotus* | *Fagus sylvatica* | CLC | ECM | 1 |
| Pilcr1 | *Piloderma croeus* | *Quercus robur* | CLC | ECM | 1 |
| Oidma1 | *Oidiodendron maius* | *Vaccinium myrtillus* | CLC | ERM | 1 |
| Sebve1 | *Sebacina vermifera* | *Arabidopsis thaliana* | CLC | ORM | 1 |
| Tulca1 | *Tulasnella calosporra* | *Serapias vomeracea* | CLC | ORM | 1 |
| Melbi2 | *Meliniomyces bicolor* | *Vaccinium myrtillus* | CLC | ERM | 2 |
| Rhier1 | *Rhizoscyphus ericaceae* | *Vaccinium myrtillus* | CLC | ERM | 2 |
| Tubma1 | *Tuber magnatum* | *Quercus robur* | CLC | ECM | 3 |
| Cenge3 | *Cenococcum geophilum* | *Pinus sylvestris* | CLC | ECM | 4 |
| Acema1 | *Acephala macrosclerotiorum* | *Pinus sylvestris* | HISAT/ DESeq2 | ECM | 5 |
| Lacbi2 | *Laccaria bicolor* | *Populus tremula x alba* | HISAT/DESeq2 | ECM | 6 |
| Trima3 | *Tricholoma matsutake* | *Pinus sylvestris* | CLC | ECM | 5 |

**Table S2: Lectin content in the predicted proteomes of the Agaricomycetes fungal class sorted by ecological niche.**

| | LysM-like | Jacalin-like | Natterin-like | Monocot-lectin like | Fungal prism lectins | AAL-like PropLec6A | PLL-like PropLec7A | PVL-like PropLec7B | Tectonin PropLec6B | galectin-like | Physarum lectin | Yeast Emp L-type | cyanovirin-like | Fungal fruit body lectin | Oomycete cytolysin | CBM67-like | H-type lectin | Ig-like | Boletus and Laetiporus b- | Clitocybe lectin-like | Coprinus b-trefoil lectin | Earthworm lectin | Fungi and Clostridium b- | Ricin-like | Ginkbilobin-like | hevein-like | Lyophyllum ginkbilobin-like | | |
|---|---|---|---|---|---|---|---|---|---|---|---|---|---|---|---|---|---|---|---|---|---|---|---|---|---|---|---|---|---|
| **Endophyte** | | | | | | | | | | | | | | | | | | | | | | | | | | | | | classes |
| *Piriformospora indica* | 5 | | | | | | | | | | | | | 3 | | | | | | | | | | | | | | 8 | 2 |
| **Orchid mycorrhizae** | | | | | | | | | | | | | | | | | | | | | | | | | | | | | 0 |
| *Sebacina vermifera* | 1 | | | | | | | | | 1 | | 1 | 2 | | | | | | | 2 | | | | | | | | 7 | 5 |
| *Tulasnella calospora* | | | | | | 1 | | | | | | | 1 | | | | 6 | | | | 3 | 1 | 1 | | | | | 13 | 6 |
| **Ectomycorrhizae** | | | | | | | | | | | | | | | | | | | | | | | | | | | | | 0 |
| *Amanita muscaria* | | | | | | | | | | | | | | | | 5 | 3 | 3 | | | | 1 | | | | | | 12 | 4 |
| *Amanita rubescens* | | | | | | | | 3 | 6 | | | | | 1 | | 11 | 6 | | | 4 | 1 | | | | | | | 32 | 7 |
| *Boletus edulis* | | | | | | | | | | | | | | 9 | | | 3 | | 3 | | 5 | | | | | | | 20 | 4 |
| *Cantharellus anzutake* | | | | | | | | | 2 | | | | 1 | | | | | | | | | | | | | | | 3 | 2 |
| *Cortinarius glaucopus* | | | | | | 4 | | 5 | 4 | | | 1 | | | | | 4 | | 1 | | | | | 1 | | 1 | | 21 | 8 |
| *Gautieria morchelliformis* | | | | | | | 6 | | | | | 1 | | 1 | | | 2 | | | | | | | | | | | 10 | 4 |
| *Gyrodon lividus* | | | | | | | | | | | | | 1 | 8 | | | 3 | | | | | | | | | | | 12 | 3 |
| *Hebeloma cylindrosporum* | | | | | | | 1 | 3 | 10 | 1 | | | 1 | 2 | | | | | | | | | | | 2 | | | 20 | 7 |
| *Hydnum rufescens* | | | | | | | | | | | | | | | | | | | | | 3 | | | | | | | 3 | 1 |
| *Hysterangium stoloniferum* | | | | | | 1 | | | | | | 1 | | | | | 3 | | | | | | | | | | | 5 | 3 |
| *Laccaria amethystina* | | | | | | | | 3 | 5 | 5 | 2 | | | 1 | | | | | | | 4 | | | | | | | 20 | 6 |
| *Laccaria bicolor* | | | | | | | | 1 | 3 | 2 | 2 | | 1 | 1 | | | | | 1 | | 3 | | | | | | | 14 | 8 |
| *Lactarius quietus* | | | | | | | | | | | | 1 | | 4 | | | 11 | | | | | | | | | | | 16 | 3 |
| *Melanogaster broomeianus* | | | | | | | | | | 2 | | | | 2 | | | | | | | | | | | | | | 4 | 2 |
| *Paxillus adelphus* | | | | | | | | | 1 | | | | | 3 | | | | | | | | | | | | | | 4 | 2 |
| *Paxillus ammoniavirescens* | | | | | | | | | | | | | | 5 | | | 3 | | | | | | | | | | | 8 | 2 |
| *Paxillus involutus* | | | | | | | | | | | 5 | | | 8 | | | 4 | | | | | | | | | | | 17 | 3 |
| *Piloderma croceum* | | | | | | | | | | | | 1 | 1 | | | | | | | | | | 1 | | | | | 3 | 3 |
| *Pisolithus tinctorius* | | | | | | | | | | | | | 4 | | | | | | | | | | | | | | | 4 | 1 |
| *Rhizopogon vesiculosus* | | | | | | 1 | | | | | | | | 1 | | | | | | 4 | 4 | | | | | | | 10 | 4 |
| *Rhizopogon vinicolor* | | | | | | 1 | | | | | | | | 1 | | | | | | 4 | 4 | | | | | | | 10 | 4 |
| *Russula ochroleuca* | | | | | | | | | | | | | | 2 | | | 2 | | | | | | | | | | | 4 | 2 |
| *Scleroderma citrinum* | | | | | | | | | | | | | 20 | | | | | | | | | | | | | | | 20 | 1 |
| *Suillus brevipes* | | | | | | 2 | | | | | | | | 1 | | | | | | 1 | | | | | | | | 4 | 3 |
| *Suillus luteus* | | | | | | 2 | | | | | | | | 1 | | | | | | | | | | | | | | 3 | 2 |
| *Thelephora ganbajun* | | | | | | | | | | | | | | | | | | | 4 | | | | | | | | | 4 | 1 |
| *Thelephora terrestris* | | | | | | 5 | | | | | | | | 1 | | | | | | | | | | | | | | 6 | 2 |
| *Tricholoma matsutake* | | | | | | | | | | 1 | | | | 1 | | | | | 1 | | | | | | | | | 3 | 3 |
| *Xerocomus badius/Imleria badia* | | | | | | | | | 2 | 2 | | | 1 | 2 | | | 2 | | 1 | 1 | 8 | | | | | | | 19 | 8 |
| **Grass decayer** | | | | | | | | | | | | | | | | | | | | | | | | | | | | | 0 |
| *Agaricus bisporus* | | | | | | | | | | | | | | 1 | | | | | | | | | | 2 | 2 | | | 5 | 3 |
| *Amanita thiersii* | | | | | | | | | | 1 | | 1 | | | | 2 | | | | | | | | | | | 6 | 10 | 4 |
| **Litter decayer** | | | | | | | | | | | | | | | | | | | | | | | | | | | | | 0 |
| *Ceriporiopsis (Gelatoporia)* | | | 4 | | | | | | | | | 1 | | | | | | 1 | | | | | | | | | | 6 | 3 |
| *Coprinellus micaceus* | | | | | | | | 11 | | | | | 11 | | | | 11 | | 1 | | | | | 3 | 2 | 2 | | 41 | 7 |
| *Coprinopsis cinerea* | | | | | | | | 2 | | 6 | | | | | | | 5 | | | 4 | 4 | | | 4 | | | | 25 | 6 |
| *Coprinopsis marcescibilis* | | | | | | | | 3 | | 6 | | 1 | | | | | | | | | | | | 3 | 5 | | | 18 | 5 |
| *Fibulorhizoctonia sp.* | | | | | | | | | | | | | 2 | 20 | | | 7 | | 1 | | 1 | | | 1 | | | | 32 | 6 |

| | | | | | | | | | | | | | | | | | | | | | | | | | | | | | |
|---|---|---|---|---|---|---|---|---|---|---|---|---|---|---|---|---|---|---|---|---|---|---|---|---|---|---|---|---|---|
| *Gymnopus androsaceus* | 1 | | | | | 2 | | | | | | | 5 | 1 | | 1 | 3 | | | | | | 1 | | 6 | | | 20 | 8 |
| *Gymnopus luxurians* | | | | 7 | | | | | | | | 1 | 10 | 3 | | 6 | | | | | | 1 | | 1 | 4 | | | 33 | 8 |
| *Leucoagaricus gongylophorus* | | | | | | | | | | | | | | | | | | | | | | | | 1 | | | | 1 | 1 |
| *Marasmius fiardii* | | | | 2 | | 3 | | | 5 | | | | | 1 | | | | | | 4 | | | | 3 | | | | 18 | 6 |
| *Mycena galopus* | 5 | | | | | 5 | | | | 2 | | | 19 | | 1 | 16 | 2 | 1 | 1 | 2 | 1 | | | 3 | 8 | 5 | | 71 | 14 |
| *Polyporus arcularius* | | | | | | | | | | | | | | | 1 | | | 2 | | | | | 1 | | | 1 | | 5 | 4 |
| *Psilocybe serbica* | | | | | 2 | 2 | | | 2 | | | | 4 | 4 | | 2 | 1 | 1 | 1 | | 3 | | 1 | | 4 | 1 | | 28 | 13 |
| *Pterula gracilis* | | | | | | 1 | | | | | | 1 | | 2 | 1 | 1 | | | 1 | | | | | | | | | 7 | 6 |
| *Sphaerobolus stellatus* | | | | | | 2 | 7 | | | | | | 9 | 4 | | 1 | | | 1 | | 8 | | 3 | 1 | 3 | 1 | | 40 | 11 |
| **Generalist decayer** | | | | | | | | | | | | | | | | | | | | | | | | | | | | | 0 |
| *Pleurotus ostreatus* | | 2 | | | | | | | | | | | 3 | 2 | | 4 | | | | | 1 | | 1 | 1 | | | | 14 | 7 |
| *Rhodosporidium toruloides* | | | | | | | | | | | | 1 | | | | | | | | | | | | | | | | 1 | 1 |
| *Serpula lacrymans* | | | | | | 1 | | | | | | | 2 | | | | | | | | 1 | | 3 | | | | | 7 | 4 |
| **Wood decayer** | | | | | | | | | | | | | | | | | | | | | | | | | | | | | 0 |
| *Auricularia subglabra* | | | | | | | | | | | | | 17 | 4 | | | 5 | 1 | | 1 | | 1 | | | | | 1 | 30 | 7 |
| *Auriculariopsis ampla* | | | | | | 1 | | | | | | 1 | | | | | | | | | | | | 3 | | | | 5 | 3 |
| *Bjerkandera adusta* | | | | | | 6 | | | | | | 1 | 1 | 7 | | | 2 | | | 1 | | | 1 | | | | | 19 | 7 |
| *Clavulina sp./Sistotrema* | | | | | | | | | | | | | 1 | 1 | | | | | | 1 | | | 2 | | | | | 5 | 4 |
| *Coniophora olivacea* | | | | | | | | | | | | 1 | 2 | 2 | | | | | | | 1 | | | | | | | 6 | 4 |
| *Coniophora puteana* | | | | | | | | | | | | 1 | 1 | | | | | | | | 1 | | | | | | | 3 | 3 |
| *Crucibulum laeve* | | | | | | | 1 | | | | | | 1 | 3 | | 3 | | | | 1 | | | | 2 | 9 | | | 20 | 7 |
| *Cylindrobasidium torrendii* | | | | 1 | | | | | | | | | | 1 | | | | | | | | | | | | | | 2 | 2 |
| *Dentipellis sp.* | | | | | | | | | | | | | 1 | 3 | | | 3 | | | 1 | | | | | | | | 8 | 4 |
| *Dichomitus squalens* | | | 3 | | | | | | | | | 1 | 1 | | | | | 3 | | 4 | 1 | | | | | | | 13 | 6 |
| *Exidia glandulosa* | | | | | | | | | | | | | 20 | 2 | 1 | | 1 | | | 1 | | | | | 3 | 3 | | 31 | 7 |
| *Fibroporia radiculosa* | | | | | | | | | | | | | 1 | | | | | | 1 | | | | | | | 1 | | 3 | 3 |
| *Fistulina hepatica* | | | | | | | | | | | | 1 | | | | | | | | | | | | | | | | 1 | 1 |
| *Fomitopsis pinicola* | | | | | | | | | | | | 1 | | | | | | | 1 | 2 | | | | | | | | 4 | 3 |
| *Galerina marginata* | | | | | | | | 2 | 2 | 1 | | | 6 | 1 | | 7 | | | 1 | | | | | 1 | 6 | | 1 | 28 | 10 |
| *Ganoderma sp.* | | | | | | | | | | | | 1 | | | 1 | | 1 | | | | | | | | | | | 3 | 3 |
| *Gloeophyllum trabeum* | | | | | | | | | | | | | 2 | | | | | | | | | | | | | | | 2 | 1 |
| *Heliocybe sulcata* | | | | | | | | | | | | 1 | 1 | | | | | | | | | | | | | | | 2 | 2 |
| *Hypholoma sublateritium* | | | | | 1 | | | 1 | | | | | 5 | 1 | | 3 | 1 | 1 | | | | | | 1 | | | | 14 | 8 |
| *Lentinellus vulpinus* | | | | 1 | | | | | | | | | 1 | | | | | 1 | | 1 | | | 3 | | | | | 7 | 5 |
| *Lentinula edodes* | | 1 | | | | | | | | | | | | | | | | | | | | | | | 1 | | | 2 | 2 |
| *Lentinus tigrinus* | | | | | | | | | | | | | | | 1 | | | 2 | | | | | 1 | | | | | 4 | 3 |
| *Neolentinus lepideus* | | | | | | | | | | | | 1 | | | | | | | | | | | | | | | | 1 | 1 |
| *Obba rivulosa* | | | 4 | 1 | | | | | | | | 1 | | | | | | 2 | | | | | | | | | | 8 | 4 |
| *Omphalotus olearius* | | | | | | | | | | | | | | | | 1 | | 1 | | | | | | | | | | 2 | 2 |
| *Peniophora sp.* | | | | 1 | | | | | | | | 2 | | | | | | | 2 | | | | 5 | | | 2 | | 12 | 5 |
| *Phanerochaete carnosa* | | | | | | 2 | | | | | | | | 2 | | | 6 | | | | | | | | | | | 10 | 3 |
| *Phanerochaete chrysosporium* | | | | | | 6 | | | | | | | 1 | 2 | | | 1 | | | | | | | | | | | 10 | 4 |
| *Phlebia brevispora* | | | | | | 1 | | | | | | | 1 | 1 | | | 2 | 2 | | 1 | | | | 2 | 7 | 2 | | 19 | 9 |
| *Phlebia centrifuga* | | | | | | | | | | | | | 3 | 1 | | | 6 | | | | | | | | | | | 10 | 3 |
| *Phlebia radiata* | | | | | | 3 | | | | 1 | | | | 1 | | | 8 | | | 1 | | | | 7 | 9 | 2 | | 32 | 8 |
| *Phlebiopsis gigantea* | | | | | | 4 | | | | | | | 1 | 1 | | | 2 | | | 1 | | | | | | | | 9 | 5 |
| *Plicaturopsis crispa* | | | | | | | | | | | | 1 | | | | | | | | | | | 1 | | | | | 2 | 2 |
| *Pluteus cervinus* | | 1 | | | | 38 | | | | 1 | | | | 1 | | 1 | | 1 | | | | | | 3 | | 2 | | 48 | 8 |
| *Polyporus brumalis* | | | | | | | | | | | | | | | 1 | | | 1 | | | | | 1 | | | 1 | | 4 | 4 |
| *Postia placenta* | | 1 | | | | | | | | | | 1 | 3 | 4 | | | 2 | 2 | | 7 | 3 | | | | | | | 23 | 8 |
| *Punctularia strigosozonata* | | | | | | | | | | | | | | 3 | | | 1 | | | | | | | | | | | 4 | 2 |
| *Pycnoporus cinnabarinus* | | | | | | | | | | | | 1 | | | | | | | | | | | 1 | | | | | 2 | 2 |
| *Ramaria rubella* | | | 1 | | | 1 | 3 | | | | | | | | | | | | | | | | | 2 | | | | 7 | 4 |

| | | | | | | | | | | | | | | | | | | | | | | | | | | | | | |
|---|---|---|---|---|---|---|---|---|---|---|---|---|---|---|---|---|---|---|---|---|---|---|---|---|---|---|---|---|---|
| *Schizophyllum commune* | | | | 1 | | 2 | | | | | | 1 | | | | | | | | | | | | 3 | | | | 7 | 4 |
| *Schizopora paradoxa* | | | | 2 | | | | | | | | 1 | 6 | 1 | | | | 1 | | 1 | | | | | | 1 | | 13 | 7 |
| *Serpula himantioides* | | | | | | 1 | | | | | | | 3 | | | | | | | | 3 | | 1 | | | | | 8 | 4 |
| *Stereum hirsutum* | | | | | | | | | | | | 1 | | 1 | | | 1 | | | | | | | | | | | 3 | 3 |
| *Trametes pubescens* | | | | | | | | | | | | | | | | | | 2 | | | | | | | | | 2 | 4 | 2 |
| *Trametes versicolor* | | | | | | | | | | | | | | | | | | 1 | | | | | | | | | 2 | 3 | 2 |
| **Plant pathogen/Wood decayer** | | | | | | | | | | | | | | | | | | | | | | | | | | | | | 0 |
| *Armillaria cepistipes* | | | | | | 2 | | | | | | | | | | 16 | | | 1 | | 4 | | 2 | 1 | | | | 26 | 6 |
| *Armillaria gallica* | | | | | | 3 | | | | | | | | | | 16 | | | 1 | | 3 | | 3 | 1 | | | | 27 | 6 |
| *Fomitiporia mediterranea* | | | | | | | | | | | | 1 | | | | | | | | 2 | | | | | | | | 3 | 2 |
| *Laetiporus sulphureus* | | | | | | 1 | | | | | | | 1 | 1 | | | | | 2 | | | | | | | | | 5 | 4 |
| **Plant pathogen** | | | | | | | | | | | | | | | | | | | | | | | | | | | | | 0 |
| *Armillaria mellea* | | | 1 | | | 1 | | | | | | | | | | 1 | | | 3 | | 5 | | 2 | 1 | | | | 14 | 7 |
| *Armillaria ostoyae* | | | | | | 2 | | | | | | | | | | 16 | | | 2 | | 3 | | 2 | | | | | 25 | 5 |
| *Armillaria solidipes* | | 1 | | | | 2 | | | | | | | | | | 17 | | | 1 | | 4 | | 1 | | | | | 26 | 6 |
| *Heterobasidion annosum* | | | | | | | | | | 1 | | | 2 | | | | | | | | | | | | | | | 3 | 2 |
| *Moniliophthora perniciosa* | | 1 | | | | | | | | | | | | | 3 | | | | | | | | | | | | | 4 | 2 |
| *Rhizoctonia solani* | | 1 | | | | | | | | | | | 8 | | | | 3 | | | 4 | 3 | 3 | | | | | | 22 | 6 |
| *Wolfiporia cocos* | | | | | | | | | | | | | | 1 | | | 1 | | | | | | | | | | | 2 | 2 |
| **Overall total** | **12** | **8** | **13** | **16** | **3** | **109** | **18** | **34** | **42** | **33** | **9** | **32** | **188** | **141** | **10** | **130** | **127** | **29** | **32** | **56** | **85** | **7** | **38** | **51** | **71** | **25** | **12** | **1331** | |
| **Number of species** | **4** | **7** | **5** | **8** | **2** | **32** | **5** | **10** | **11** | **15** | **3** | **31** | **44** | **53** | **8** | **20** | **36** | **19** | **22** | **25** | **27** | **5** | **22** | **24** | **15** | **14** | **5** | **107** | |
| % species | 4% | 7% | 5% | 7% | 2% | 30% | 5% | 9% | 10% | 14% | 3% | 29% | 41% | 50% | 7% | 19% | 34% | 18% | 21% | 23% | 25% | 5% | 21% | 22% | 14% | 13% | 5% | 100% | |

**Table S3 : Details of lectins identified in the genome of *Laccaria bicolor***

| **Lectin class** | **#** | **Mycocsm AC (score)** | **NCBI AC** | **Protein name** |
|---|---|---|---|---|
| Tectonin PropLec6B | 3 | Lacbi2:399271 (0.73)<br>Lacbi2:399270 (0.67)<br>Lacbi2:322629 (0.27) | XP_001876432.1 | tectonin 2 |
| Coprinus β-trefoil lectin | 3 | Lacbi2:330799 (0.48)<br>Lacbi2:327918 (0.42)<br>Lacbi2:691792 (0.25) | XP_001885184.1 | predicted protein [*Laccaria bicolor* S238N-H82] |
| Galectin like | 2 | Lacbi2:236913 (0.36)<br>Lacbi2:312069 (0.35) | XP_001883510.1 | galectin [*Laccaria bicolor* S238N-H82] |
| Physarium lectin | 2 | Lacbi2:381649 (0.39)<br>Lacbi2:322629 (0.39) | XP_001875654.1 | ricin-containing lipase tectonin-like |
| Oyster lectin | 2 | Lacbi2:585014 (0.28)<br>Lacbi2:448672 (0.27) | XP_001880964.1 | predicted protein [*Laccaria bicolor* S238N-H82] |
| Fungal fruit body lectin | 1 | Lacbi2:185716 (0.54) | XP_001885326.1 | predicted protein, partial |
| Boletus and Laetiporus β-trefoil lectin | 1 | Lacbi2:318163 (0.30) | XP_001879265.1 | predicted protein [*Laccaria bicolor* S238N-H82] |
| Cyanovirin like | 1 | Lacbi2:327824 (0.42) | XP_001881773.1 | predicted protein [*Laccaria bicolor* S238N-H82] |
| P domain of calnexin and reticulin | 1 | Lacbi2:399410 (0.50) | XP_001874124.1 | calnexin [*Laccaria bicolor* S238N-H82] |
| Ergic vip L type | 1 | Lacbi2:399414 (0.48) | XP_001888824.1 | ERGIC53, mannose lectin |
| P-type lectin like | 1 | Lacbi2:642707 (0.29) | XP_001874815.1 | predicted protein [*Laccaria bicolor* S238N-H82] |
| PVL like PropLec7B | 1 | Lacbi2:692684 (0.67) | XP_001891161.1 | predicted protein, partial |
